# Bayesian inference by visuomotor neurons in prefrontal cortex

**DOI:** 10.1101/2024.09.23.614567

**Authors:** Thomas Langlois, Julie A. Charlton, Robbe L. T. Goris

## Abstract

Perceptual judgements of the environment emerge from the concerted activity of neural populations in decision-making areas downstream of sensory cortex [1, 2, 3]. When the sensory input is ambiguous, perceptual judgements can be biased by prior expectations shaped by environmental regularities [4, 5, 6, 7, 8, 9, 10, 11]. These effects are examples of Bayesian inference, a reasoning method in which prior knowledge is leveraged to optimize uncertain decisions [12, 13]. However, it is not known how decision-making circuits combine sensory signals and prior expectations to form a perceptual decision. Here, we study neural population activity in the prefrontal cortex of macaque monkeys trained to report perceptual judgments of ambiguous visual stimuli under two different stimulus distributions. We analyze the component of the neural population response that represents the formation of the perceptual decision (the decision variable, DV), and find that its dynamical evolution reflects the integration of sensory signals and prior expectations. Prior expectations impact the DV’s trajectory both before and during stimulus presentation such that DV trajectories with a smaller dynamic range result in more biased and less sensitive perceptual decisions. These results reveal a mechanism by which prefrontal circuits can execute Bayesian inference.

---

Perceptual systems infer properties of the environment from sensory measurements that can be ambiguous. However, prior knowledge can be leveraged to disambiguate the interpretation [13]. This inference strategy typically manifests as perceptual interpretations that are biased towards the prior expectation. Such biases may reflect implicit knowledge of statistical regularities that are stable features of the environment, such as the tendencies of sunlight to come from above [4], image velocities to be slow [6, 7], and cardinal orientations to be over-represented in visual scenes [14]. However, biased perceptual interpretations can also reflect knowledge of statistical regularities that are context-specific and short-lived in nature [8, 9, 10, 11]. The diversity of experimental settings under which prior-induced perceptual biases occur suggests that a general neural mechanism may underlie these effects [15, 16, 17].

Prior expectations about sensory stimulation modulate neural activity in decision circuits in various ways. Context cues that signal specific environmental statistics can selectively modulate activity of single cells before stimulus onset [18], during stimulus presentation [19, 20, 21, 22], and while the perceptually-driven behavior is being produced [21, 22]. At the population level, these effects conspire to bias neural representations towards the prior expectation [22]. Because these neural effects co-occur with biases in perceptual reports, they are thought to reflect the neural computations that govern the perceptual interpretation of the environment [18, 20, 21, 22]. However, there is an alternative explanation that cannot be ruled out. In all of these previous studies, perceptual interpretations had a fixed relationship to the overt motor response. Biases in perceptual reports thus coincided with biases in motor responses. Decision-making circuits are typically involved in action planning [23, 24, 3]. It is thus not clear whether the reported neural correlates of perceptual expectation pertain to perceptual inference, motor planning, or a mixture of the two.

To obtain an unobstructed view on the neural correlates of perceptual expectation and inference, we used a task that requires flexible reporting of perceptual decisions [3]. Monkeys judged whether a visual stimulus was oriented clockwise or counter-clockwise from vertical and communicated their decision with a saccadic eye movement towards one of two visual targets (Fig. 1a). The meaning of each response option was signaled by the target’s orientation (clockwise vs counterclockwise) and was unrelated to its spatial position (one target was placed in the neurons’ estimated motor response field, the other was placed on the opposite side of the fixation mark, see Methods). Because the spatial configuration of the choice targets varied randomly from trial to trial, changes in prior stimulus statistics biased the animals’ perceptual reports, but not their overt motor responses. While the animals performed this task, we recorded extracellular responses from neural ensembles in the pre-arcuate gyrus (Supplemental Fig. 1), an area of prefrontal cortex (PFC) involved in the selection of saccadic eye movements [25, 26] that represents visuomotor deliberation [27, 28]. We previously found that the population activity initially represents the formation of a perceptual choice, before transitioning into the representation of the motor plan [3]. Here, we study how prior expectations and sensory signals shape the perceptual decision-making process at the single trial level. Our results reveal that prior expectations selectively change the dynamic evolution of the decision-related component of population activity in PFC. Expectations impact the starting point, slope, and dynamic range of the stimulus-driven trajectory in neural activity, culminating in more biased and less sensitive perceptual decisions.

**Figure 1.**
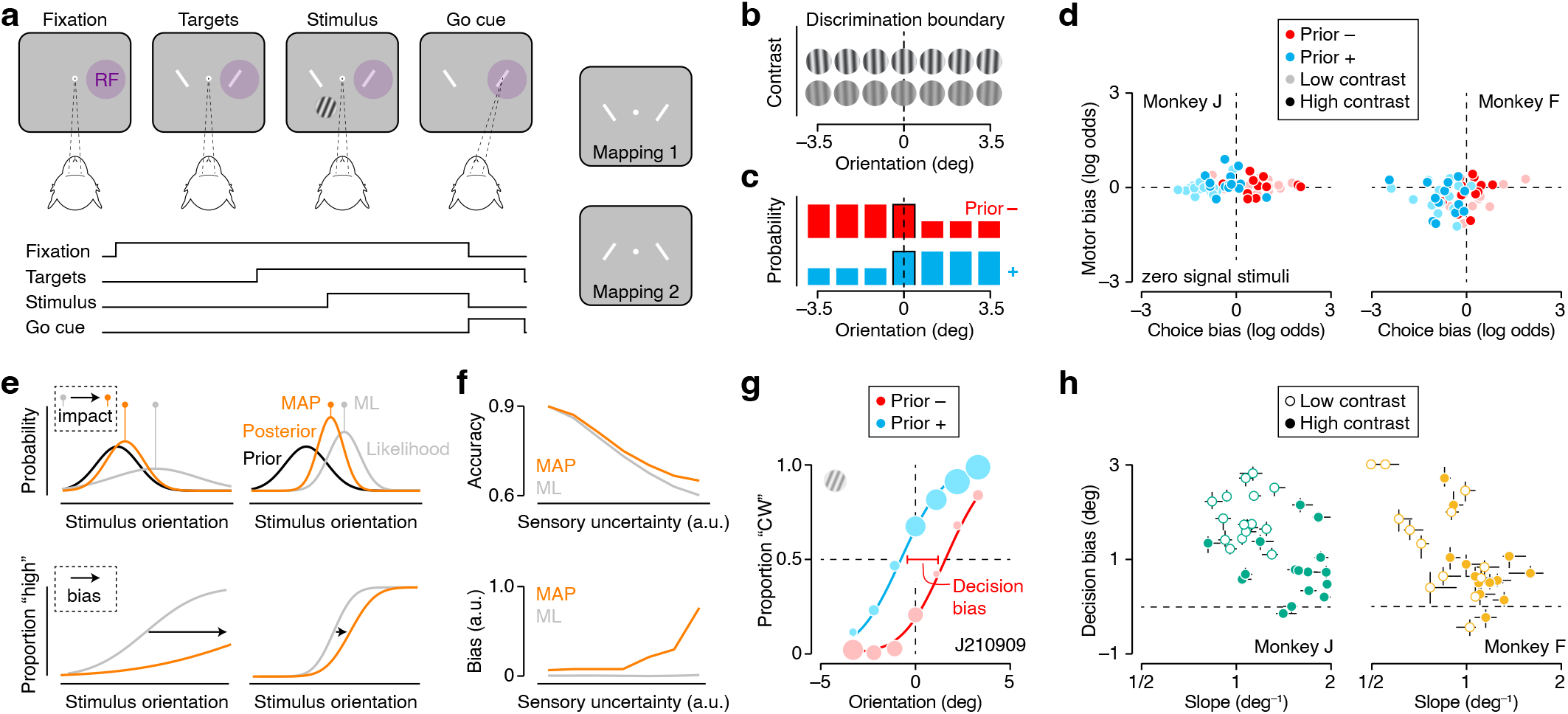
Flexible orientation discrimination under different prior statistics: task and behavior. (**a**) Orientation discrimination task, task sequence. Each trial begins when the observer initiates fixation. The shape of the fixation mark indicates the prior distribution from which the stimulus will be drawn. After the observer fixates for 500 ms, two choice targets appear, followed by the stimulus. The observer judges whether the stimulus is rotated clockwise or counterclockwise relative to vertical and communicates this decision with a saccade towards the matching choice target. Correct decisions are followed by a juice reward. One of the choice targets is placed in the neurons’ presumed motor response field (see Methods). The spatial organization of the choice targets varies randomly from trial-to-trial, giving rise to two stimulus-response mapping rules. (**b**) Stimuli varied in orientation and contrast. (**c**) Stimuli were drawn from one of two orientation distributions. (**d**) Comparison of choice and motor bias for the randomly rewarded ambiguous stimuli (orientation = 0 deg) under both priors for both monkeys (see Methods). (**e**) An ideal Bayesian decision-maker evaluates the likelihood of every possible state of the sensory environment and multiplies this distribution with the prior probability of encountering each state to obtain a posterior belief function. The posterior informs the decision. More ambiguous sensory measurements yield a broader likelihood function, and ultimately more biased decisions. (**f**) Choice accuracy (top) and bias (bottom) in our task under a maximum likelihood (ML) and maximum posterior probability (MAP) inference strategy (see Methods). The performance benefit conferred by accurate prior knowledge grows with sensory uncertainty (top), as does the magnitude of the decision bias (bottom). (**g**) Psychophysical performance for monkey J in an example recording session. Proportion ‘clockwise’ choices for low contrast stimuli is shown as a function of stimulus orientation under both priors. Symbol size reflects number of trials (total: 1,875 trials). The curves are fits of a behavioral model (see Methods). (**h**) Decision bias plotted as a function of orientation sensitivity for both monkeys (left: Monkey J, right: Monkey F). Each symbol summarizes data from a single recording session. Closed symbols: high contrast stimuli, open symbols: low contrast stimuli. Error bars reflect the IQR of the estimate.

## Results

### Task and Behavior

Two rhesus macaques were trained to perform a visual orientation discrimination task in which they communicated perceptual decisions under two different stimulus-response mapping rules (Fig. 1a). The data were previously described in detail in ref. [3]. Task difficulty was controlled by manipulating stimulus orientation and contrast (Fig. 1b). Monkeys received a reward if they selected the appropriately oriented choice target. They performed the task similarly well under both mapping rules (median performance: monkey F = 79.7% correct; monkey J = 79.6% correct; median difference in performance across mapping rules: monkey F = 2.58%, *P* = 0.002; monkey J = 0.5%, *P* = 0.57; Wilcoxon signed-rank test). We manipulated the animals’ prior expectations by sampling stimulus orientation from one of two skewed distributions (Fig. 1c). On each trial, the shape of the fixation mark signaled the current prior condition. The prior condition and stimulus contrast were switched across short blocks of trials (see Methods). The prior manipulation selectively biased the animals’ perceptual judgments of ambiguous stimuli (median difference in choice bias for vertical gratings: monkey F = 1.22 log odds, *P <* 0.001; monkey J = 1.49 log odds, *P <* 0.001), but not the motor responses used to communicate these decisions (median difference in motor bias: monkey F = 0.03 log odds, *P* = 0.31; monkey J = 0.03 log odds, *P* = 0.06; Fig. 1d). Together, these results suggest that our paradigm engages brain mechanisms that specifically bias perceptual judgements of ambiguous sensory inputs.

What is the nature of the inference process that underlies these biased judgements of visual stimuli? A prominent hypothesis is that these biases naturally arise under an inference strategy that seeks to make the best possible decision given ambiguous sensory measurements and prior experience [12, 13]. This notion is formalized in the framework of Bayesian inference. An ideal Bayesian decision-maker computes the posterior evidence in support of each response option by multiplying the prior probability of each stimulus orientation with the likelihood that a specific stimulus orientation gave rise to the current sensory measurement (Fig. 1e, top). As a consequence, perceptual decisions are biased for all stimulus orientations, manifesting as a horizontal shift of the psychometric function (Fig. 1e, bottom, grey vs orange line). The impact of the prior on the decision depends on the ambiguity of the sensory response. The more ambiguous the sensory response, the broader the likelihood function, and the larger the impact of the prior on the posterior (Fig. 1e, bottom left vs right panel). A Bayesian inference strategy maximizes choice accuracy, even though it results in decision biases and systematic errors (Fig. 1f, grey vs orange lines). Does this strategy explain why the monkeys’ judgments of ambiguous stimuli were biased? In keeping with this hypothesis, the monkeys’ decisions were biased for all stimulus orientations, not just vertical gratings (Fig. 1g, red vs blue symbols). Moreover, they tended to make more biased decisions in task conditions associated with higher sensory uncertainty. We estimated decision bias by measuring the separation between the prior-conditioned psychometric functions, and sensory uncertainty by calculating the slope of the psychometric function (see Methods; Fig. 1g). For both monkeys, decision bias and sensory uncertainty were significantly correlated (Spearman rank correlation: Monkey J = –0.42, *P* = 0.017; Monkey F = –0.60, *P* = 0.0015; Fig. 1h). This data pattern suggests that the animals used a perceptual decision-making strategy that resembles Bayesian inference.

### Linking PFC population activity to perceptual inference at the single trial level

The animals’ choice behavior was variable. For most experimental conditions, the overall proportion of clockwise choices was neither zero nor one (Fig. 1g). This choice variability may either arise from cross-trial variability in sensory measurements [29, 30, 31, 32] or from fluctuations in prior expectations [11]. Given that choice variability was minimal or absent for the most extreme stimulus orientations (Fig. 1g), less likely sources are noise in the choice-response mapping process or attentional “lapses”. Because the choice variability has no obvious external origin, identifying the neural factors that determine the outcome of individual decisions ultimately requires a moment-to-moment analysis of neural population activity within single trials [33, 34].

We obtained a trial-by-trial measurement of the animals’ evolving decision state by decoding a time-varying decision variable (DV) from jointly recorded neural responses using linear discriminant analysis (see Methods). The DV indicates how well the subject’s upcoming perceptual choice can be predicted from a 50 ms bin of neural population activity. In previous work, we showed that trial-averaged DV trajectories exhibit key signatures of a decision-making process [3]. Most importantly, grouping trials by stimulus strength and choice accuracy revealed a graded representation of sensory evidence (the more the stimulus orientation deviates from vertical, the higher the sensory evidence). By contrast, the action-planning component of neural activity did not show these effects. We concluded that the decision-making process was implemented as a competition between possible perceptual interpretations, independent of the ensuing action. Here, we build on these findings to develop an analysis aimed at uncovering how decision-making circuits integrate perceptual expectations and sensory signals.

Consider the DV trajectories of two example trials, measured from the same set of neurons in the same experimental condition (Fig. 2a, symbols). Both trials yielded identical overt choice behavior (a clockwise choice indicated with a rightward saccade). The raw DV trajectories are noisy, but share some clear commonalities. On both trials, the DV value hovers around zero before stimulus onset. Following stimulus onset, the DV ramps up to a peak value, after which it decays steadily. We speculate that the peak occurs around the time of choice commitment, after which the animal starts to prepare the corresponding motor response [3]. Closer inspection of the trajectories reveals that the build-up towards the peak differs in a number of ways. The excursions seem to start from a different baseline level and the ramping phase appears to differ in slope and amplitude. We hypothesize that these features capture an important aspect of the temporal evolution of the animal’s decision state. We therefore sought to obtain quantitative estimates of these trajectory features for each trial.

**Figure 2.**
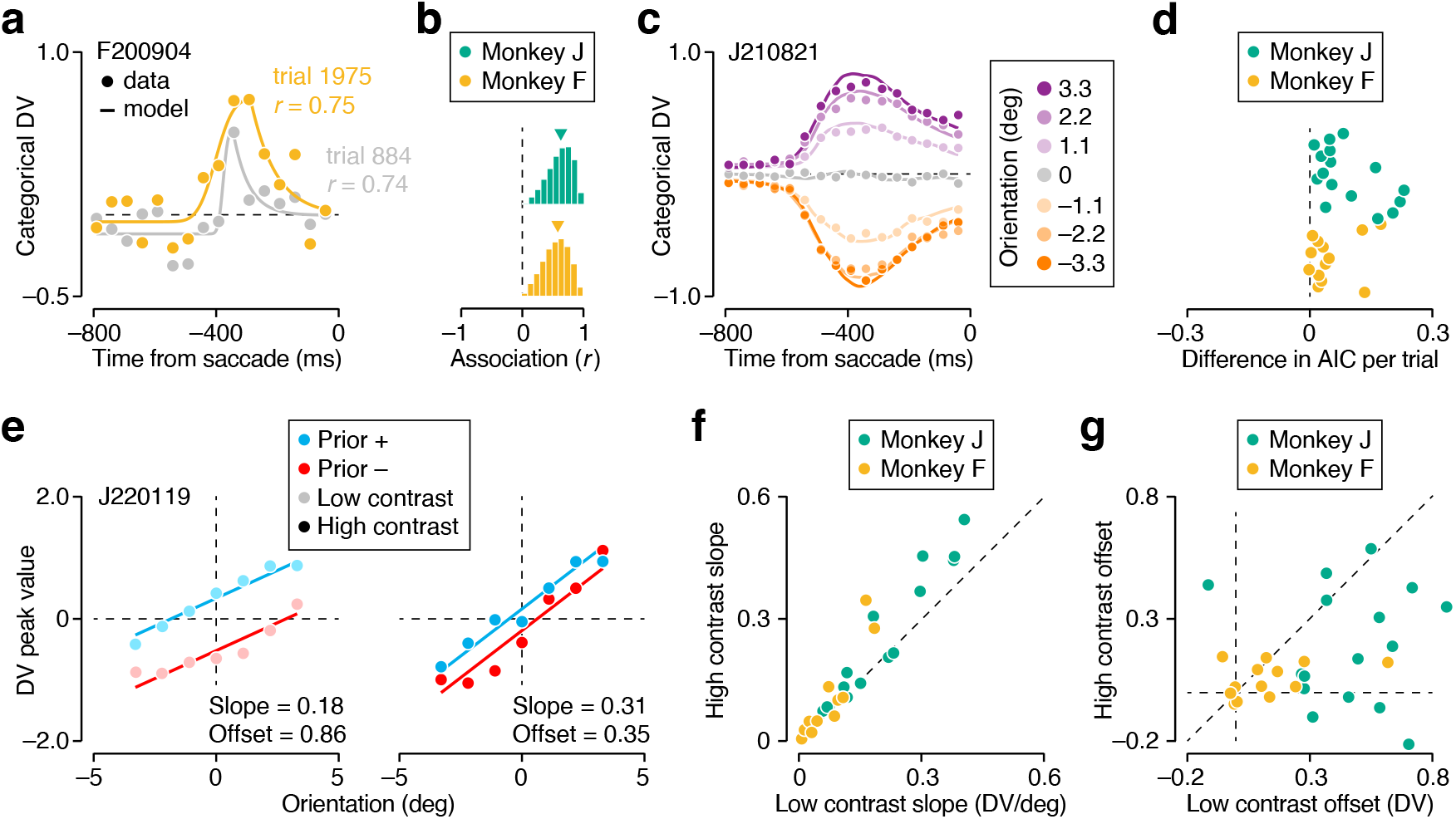
Extracting neural correlates of Bayesian-like perceptual inference at the single trial level from PFC population activity. (**a**) Temporal evolution of the categorical DV on two different trials from the same recording session with identical experimental parameters and choice behavior (prior = counter-clockwise skew, stimulus orientation = 0 deg, stimulus contrast = low, mapping rule = 1, choice = “clockwise”). Symbols show the raw DV estimates, lines the fit of a model. (**b**) Distribution of the correlation between model DV-trajectories and raw DV estimates across all recording sessions. (**c**) Trial-averaged temporal evolution of the categorical DV, split out by stimulus orientation, for a single recording session. (**d**) Difference in prediction error for two logistic regression analyses of the choice behavior, as measured by Akaike’s Information Criterion. The first analysis used stimulus prior, orientation and contrast as regressors, the second analysis additionally used the model DV-trajectory peak value. Positive values indicate that including the peak DV value improves prediction accuracy. (**e**) The peak value of the categorical DV depends on stimulus orientation (abscissa), stimulus contrast (left vs right panel), and stimulus prior (red vs blue). Symbols show the average value across all trials from a single recording session. Lines show the outcome of a linear regression analysis used to estimate the slope and offset of this relation at low (left panel) and high (right panel) stimulus contrasts. (**f**,**g**) Comparison of slope (**f**) and offset (**g**) for high vs low contrast stimuli.

We estimated key features of the DV trajectories by fitting a model in which trajectories evolve smoothly to the raw DV values (Fig. 2a, lines; see Methods). This way, we obtained a single “model DV-trajectory” for each trial. Due to the irregular nature of the raw DV trajectories, the correlation between model DV-trajectory and raw DV value was modest (median Pearson correlation: Monkey J = 0.63; Monkey F = 0.58; Fig. 2b). Nevertheless, the model captured the systematic structure of the trial-averaged data well. This was evident at the level of individual recording sessions (example shown in Fig. 2c). To test whether the model DV-trajectories afford additional insight into the animals’ decision state, we conducted a logistic regression analysis of the choice behavior. For each recording session, we first measured the association between the experimentally controlled variables (stimulus prior, orientation, and contrast) and choice outcome. We then asked whether additionally including the peak value of the model DV-trajectory as a regressor helped to better predict choice outcome (see Methods). We computed a standard measure for prediction error (Akaike’s Information Criterion, AIC) and found that including the peak value of the model DV-trajectories systematically improved prediction quality (manifesting as a difference in AIC that is larger than 0 in Fig. 2d). Together, these results suggests that for each trial, the model DV-trajectory provides a useful estimate of the DV’s baseline level and the slope and magnitude of its ramping phase.

We have argued that the monkeys’ behavior suggests that they combined prior expectations and sensory inputs in a manner that resembles Bayesian inference. We therefore hypothesized that the peak value of the neurally decoded DV should exhibit key signatures of a Bayesian posterior. To test this, we investigated how the model DV-trajectories’ peak value depended on the stimulus prior, orientation, and contrast. In the vast majority of recording sessions, the peak DV value exhibited the hypothesized structure, as can be seen for an example session (Fig. 2e). Specifically, the peak DV grows with stimulus orientation, but is biased by the stimulus prior. The impact of the sensory input is larger when stimulus contrast is high, as is evident from the difference in the slope of a linear regression line (Fig. 2e, left vs right panel, slope = 0.18 for low contrast stimuli and 0.31 for high contrast stimuli). Conversely, the impact of the prior expectation is larger when stimulus contrast is low, as is evident from the difference in the offset of the regression lines (Fig. 2e, left vs right panel, offset = 0.86 for low contrast stimuli and 0.35 for high contrast stimuli). These effects are representative of both animals’ data (median difference in slope between low and high contrast stimuli: monkey J = 0.02, *P* = 0.003; monkey F = 0.01, *P* = 0.007; one-sided Wilcoxon signed-rank test; Fig. 2f; median difference in offset: monkey J = -0.28, *P* = 0.016; monkey F = -0.04, *P* = 0.05; Fig. 2g). Thus, around the putative time of choice commitment, the neurally estimated decision-state exhibits key signatures of a Bayesian posterior. Having established this, we now turn to the question of how prior expectations impact the evolving decision-state over the course of a single trial.

### Prior expectations bias neural population trajectories

The effects of prior expectation on the evolving decision-state can be appreciated by considering neural responses to low contrast stimuli whose orientation matches the categorization boundary (vertical gratings). These stimuli yield sensory responses that are, on average, completely ambiguous. It follows that any systematic choice biases, summarized in Fig. 1d, are purely driven by the subject’s prior belief. As can be seen for an example recording session, the neural DV reflects this prior belief well before stimulus onset (Fig. 3a). At this point in time, the animal can only leverage knowledge about the skew of the stimulus distribution (i.e., whether a clockwise or counter-clockwise orientation is more likely) and the level of sensory uncertainty (i.e., whether a low or high contrast stimulus is more likely). For both animals, this contextual information biases the DV’s early value in the direction of the prior expectation, though the effect was more prominent for Monkey J than for Monkey F. This was evident from the DV traces averaged across all sessions (Fig. 3b) and from the model DV-trajectory value before stimulus onset (difference in median value for the –800 to –600 ms epoch: monkey J = 0.118, *P <* 0.001; two-sided Wilcoxon rank sum test; monkey F = 0.008, *P <* 0.001; Fig. 3c). As can also be seen from the average DV traces, the prior-induced bias is not static, but grows over time. To quantify this, we used the model DV-trajectories to compute the average DV value per recording session for an early and late temporal window (spanning the range from –800 to –600 ms and –500 to –300 ms). The first window precedes any stimulus-driven activity in PFC, while the second one overlaps with the putative time of choice commitment. Plotting the late against the early DV value reveals a relationship that is steeper than the line of unity (Fig. 3d). For both monkeys, the gradient of the first principle component was approximately 72.5 degrees, significantly steeper than 45 degrees (monkey J: 72.0 deg, *P <* 0.001, bootstrap analysis, see Methods; monkey F: 72.7 deg, *P <* 0.001).

**Figure 3.**
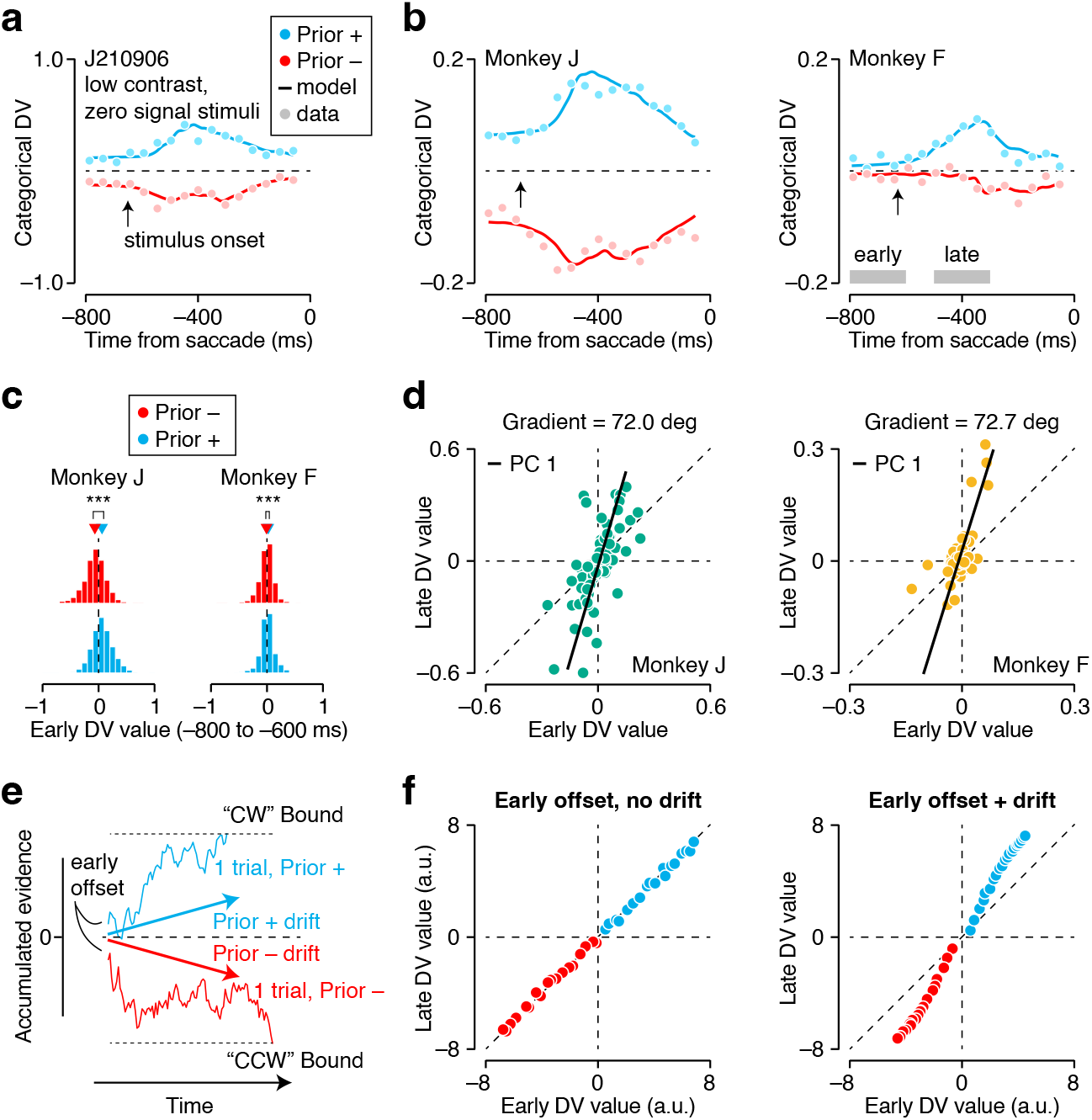
Prior expectations bias the decision variable before and during stimulus presentation. (**a**) Trial-averaged temporal evolution of the categorical DV, split by stimulus prior, for a single recording session. Only low contrast, zero signal stimuli are included. Symbols show the raw DV estimates, lines the model DV-trajectories. The average time of stimulus onset is indicated by the black arrow. (**b**) The average across all recording sessions, split by monkey (left vs right panel). (**c**) Distribution of the modelestimated early DV value across all recording sessions, split by stimulus prior (blue vs red) and monkey (left vs right). (**d**) Late DV value plotted against early DV value, split by monkey (left vs right). Each symbols summarizes the trial-averaged values of a single recording session (one point per stimulus prior). (**e**) We simulated bounded accumulation of ambiguous sensory evidence under two different prior expectations. In all simulations, the prior changes the starting point of the accumulation process (blue vs red trace, leftmost time point). In some simulations, the prior additionally changes the drift rate of the accumulation process (blue vs red arrow). (**f**) Late DV value plotted against early DV value for the simulations without (left) and with (right) prior-induced drift.

Why does the influence of the prior expectation grow during deliberation? We hypothesize that both animals integrated incoming sensory evidence over time and gave more weight to noisy orientation estimates that agreed with their prior expectation. Such a confirmation bias deviates from an ideal inference strategy, in which all sensory evidence is given the same weight, resulting in a flat average DV trace for ambiguous stimuli. However, a confirmation bias naturally arises when subjects use an inference strategy that approximates the posterior rather than computing it exactly [35]. Under approximate inference, as incoming sensory evidence is accumulated to update the posterior belief in the state of the world, the prior belief enters into the update multiple times, resulting in a positive feedback loop [35]. Critically, this explanation implies that a model comprised of an initial prior-induced offset and a bound that terminates the deliberation process does not suffice to capture the dynamically increasing influence of the prior on the DV. The extra ingredient that is needed is a prior-induced drift [35]. To test this explanation for our data, we simulated bounded accumulation of ambiguous sensory evidence and compared early and late DV values (see Methods; Fig. 3e). We varied the strength of the prior-induced effects and the height of the bound, resulting in a range of DV values (Fig. 3f). In the simulations that only included an early offset and lacked a prior-induced drift, early and late DV values were equal on average (Fig. (3f, left). We found that inclusion of a prior-induced drift was necessary to produce late DV values that were greater than the early DV values, as seen in the physiological data Fig. (3f, right). Together, these results suggest that the subjects interpreted neutral sensory evidence in a biased fashion, thus further entrenching their prior expectations.

### Relating the dynamic range of neural trajectories to decision bias and sensory uncertainty

Our analysis so far suggests that the decision-related component of neural population activity in PFC resembles a bounded accumulation process in which prior expectations influence the DV both before and during accumulation of sensory evidence. This hypothesis is in part motivated by the neural DV’s temporal evolution during presentation of ambiguous sensory stimuli (Fig. 3). To test its explanatory power, we first asked whether such a process can capture the behavioral effects of prior expectation for ambiguous as well as non-ambiguous stimuli. To this end, we extended the drift-diffusion model simulation to non-ambiguous stimulus conditions (see Methods; Extended Data Fig. 2a). In our simulation, stimulus orientation determines the mean of the momentary sensory evidence distribution, while stimulus contrast governs its spread. We further assumed that prior expectations influence the DV’s initial value and its stimulus-dependent drift in exactly the same manner across all stimulus conditions. A decision is reached when a bound is crossed, or when time is up. In the latter case, the sign of the DV determines choice outcome. Consider one specific instantiation of this process (Fig. 4a). As can be seen, prior expectations produce a horizontal shift of the psychometric functions, just like we observed in the animals’ behavior (compare with Fig. 1g). Moreover, lowering stimulus contrast makes the psychometric function more shallow and increases the decision bias (Fig. 4a, light vs dark lines), again recapitulating the animals’ behavior. Varying the model parameters can change the magnitude of the decision bias and the steepness of the psychometric functions, but it does not alter this basic pattern (Extended Data Fig. 2b).

**Figure 4.**
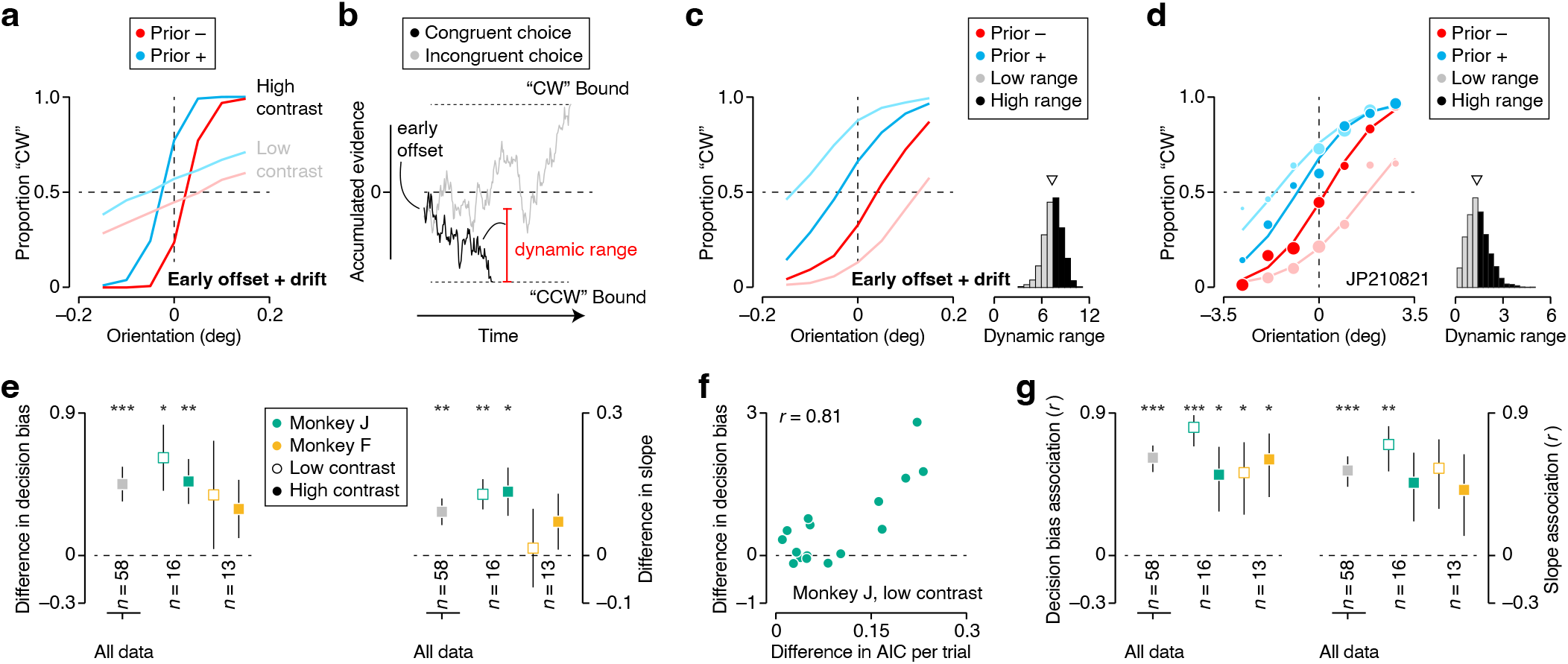
Shorter neural excursions result in more biased perceptual decisions. (**a**) We simulated bounded accumulation of ambiguous and non-ambiguous sensory evidence under two different prior expectations. In all simulations, the prior influenced the starting point and the drift rate of the accumulation process. Under this process, prior expectations induce a horizontal shift of the psychometric function (blue vs red), that is larger when the sensory evidence is less reliable (dark vs light colors). (**b**) Example of a simulated trial that resulted in a choice that was congruent with the prior expectation (black trace), and a trial that resulted in an incongruent choice (grey trace). The dynamic range of the excursion measures the distance travelled between the beginning and end of the accumulation process. (**c**) Median-split analysis of simulated choice behavior under two different prior expectations (blue vs red). Trials with a smaller dynamic range yield more biased decisions (dark vs light colors). (**d**) Median-split analysis for all low contrast trials in one example recording session. Symbol size reflects number of trials (total: 1,789 trials). (**e**) (Left) The difference in decision bias for low and high dynamic range trials across all data (grey symbols), and split out by monkey and stimulus contrast (non-grey symbols). (Right) The difference in the slope of the psychometric function for low and high dynamic range trials. Error bars show mean +-1 s.e.m. (**f**) Difference in decision bias plotted as a function of the difference in Akaike’s Information Criterion. (**g**) (Left) Association between the AIC difference and the difference in decision bias. (Right) Association between the AIC difference and the difference in the slope of the psychometric function. Error bars show *r* +-1 s.d. * *P* i 0.05, ** *P* i 0.01, *** *P* i 0.001.

The proposed decision-making process gives rise to a relationship between the dynamic range of the DV trajectory and decision bias. Specifically, simulated trials in which the eventual choice aligns with the prior expectation tend to have a smaller dynamic range (defined as the difference between the simulated DV’s peak value and its initial value), while trials in which the choice deviates from the prior expectation tend to have a larger dynamic range (Fig. 4b). This relationship arises naturally if a prior expectation influences the DV’s initial value while a fixed boundary terminates the deliberation process (or conversely, if a prior expectation influences the bounds without impacting the initial value). To examine the relationship between dynamic range and decision bias, we computed the distribution of the DV’s dynamic range across all trials within a simulated experiment and separately analyzed the choice behavior for the trials whose dynamic range was below and above the median (Fig. 4c). We observed that the effect of the prior differs substantially across the low and high dynamic range trials in the simulated dataset. Specifically, when the DV’s dynamic range is low, the decision bias is amplified and the psychometric function is more shallow (Fig. 4c, light vs dark lines). This pattern of results is specific to a decision-making process in which the prior changes the distance between the DV’s starting point and the bounds. Alternative scenarios in which the prior only induces a drift during the accumulation of sensory evidence, or in which there is no terminating boundary, do not produce a relationship between dynamic range and decision bias (Extended Data Fig. 3).

To test whether these simulated effects are present in the real data, we conducted the same median-split analysis on the neural DVs, using the model DV-trajectories to estimate each trial’s dynamic range (Methods). We discovered the same relationship between dynamic range and the animals’ decision bias. To quantify this relationship, we computed the difference between the decision bias for low and high dynamic range trials, Δ*B* = Δ*β*_*L*_ − Δ*β*_*H*_ (example shown in Fig. 4d). Across all experiments, the mean value of Δ*B* was 0.45 (*P <* 0.001, Wilcoxon signed rank test; Fig. 4e, left). Considering the data at a more granular level (per monkey and per contrast level) reveals the robustness of these results, though not every individual case reached statistical significance (Fig. 4e, left). We also measured the relationship between dynamic range and the steepness of the psychometric function, quantified as Δ*S* = Δ*σ*_*L*_ − Δ*σ*_*H*_, and again found a pattern of results consistent with the model simulation. Lower dynamic range trials tended to be associated with a shallower psychometric function (mean value of Δ*S* was 0.09, *P <* 0.01; Fig. 4e, right). For some datasets, these effects were substantial, for others they were small. Interestingly, the effects tended to be larger for datasets where the model DV-trajectories’ peak values better predicted the animal’s choice behavior (Fig. 4f). To summarize these results, we computed the correlation between Δ*B* and Δ*S* on the one hand, and the previously introduced measure for model prediction error (AIC) on the other. Across all experiments, the correlation with Δ*B* was 0.62 (*P <* 0.001; Fig. 4g, left panel), while the correlation with Δ*S* was 0.54 (*P <* 0.001; Fig. 4g, right panel). Thus, the stronger the neural correlate of the decision-making process, the more the empirical data resemble the predictions of an idealized model of how prior expectations shape this process.

We hypothesize that the neural DV’s dynamic range primarily reflects the strength of the prior expectation. Under a bounded accumulation process with a prior-induced initial offset and drift, a prior expectation reduces the dynamic range of congruent choices, but has the opposite effect on incongruent choices. In this manner, a prior expectation simultaneously amplifies the decision bias and shallows the psychometric function. Varying the strength of the prior expectation under our imagined scenario impacts both summary statistics of the observable behavior, as is evident from simulations of this process (Fig. 5a). We found that this co-occurrence was also present in our empirical observations (Fig. 5b). Datasets that exhibited a stronger association between dynamic range and decision bias (as measured by Δ*B*) also exhibited a stronger association between dynamic range and slope of the psychometric functions (correlation between Δ*B* and Δ*S*: 0.63, *P <* 0.001; Fig. 5c). This relationship remained significant after controlling for the contribution of model prediction error (AIC), a confounding variable that could have inflated this relationship (F = 10.18, *P* = 0.0025; ANCOVA with 4 levels of prediction error). Together, these results support the conclusion that shorter excursions of the neural DV yield more biased, less sensitive perceptual decisions.

**Figure 5.**
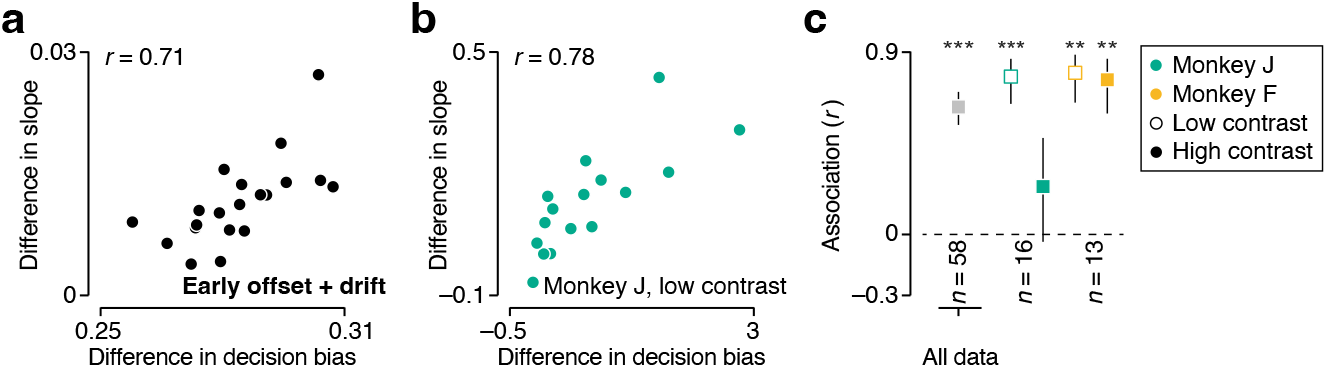
Stronger prior expectations yield decisions that are more biased and less sensitive. (**a**) We simulated bounded accumulation of sensory evidence under two different prior expectations and measured the association between the DV’s dynamic range and decision bias (abscissa) and between dynamic range and the slope of the psychometric function (ordinate). Each symbol summarizes these measurements for one simulation. We varied the strength of the prior expectation across simulations. (**b**) Difference in slope plotted against difference in decision bias for one monkey. Each symbol summarizes one recording session. Only low contrast trials were included in the analysis. (**c**) Summary of the association between both variables across data (grey symbol), and split out by monkey and stimulus contrast (non-grey symbols). Error bars show *r* +-1 s.d. * *P* i 0.05, ** *P* i 0.01, *** *P* i 0.001.

## Discussion

In this study, we investigated neural population activity in the prefrontal cortex of macaque monkeys while they judged ambiguous visual stimuli under different prior stimulus statistics. We sought to understand how decision-making circuits combine sensory signals and prior knowledge to form a perceptual decision. We used a task that requires flexible reporting of perceptual decisions and discovered that prior expectations exert numerous influences on neural population activity during decision formation. In our task, the neural correlate of decision formation resembles a tug-of-war style competition between candidate perceptual interpretations, independent of the ensuing motor response used to communicate the decision [3]. Prior expectations shift the starting point of the competition towards the more probable stimulus interpretation (Fig. 3a-c). Additionally, prior expectations selectively change the impact of each piece of sensory evidence on the competition, favoring the evidence that is congruent with the expectation (Fig. 3d-f). Finally, prior expectations shorten the distance between the starting point and end point of the decision formation trajectory (Fig. 4).

Several previous studies also used biased perceptually-driven behavior as a gateway to study the neural implementation of Bayesian computation in decision making areas [18, 19, 20, 21, 22, 36]. These studies revealed that expectations can modulate the activity of single neurons before stimulus onset [19, 18], while the stimulus is presented [20, 18, 21, 22], and while the behaviour is produced [21, 22]. However, due to the experimental design, it is impossible to say whether these effects reflect a neural correlate of perceptual inference or of motor planning. Here, we built on this work by using a task in which the perceptual and motor components of the decision process are orthogonalized (Fig. 1a). This enabled us to isolate the component of neural activity that solely pertains to the formation of a perceptual choice. Our analysis revealed population level effects that echo elements of the aforementioned findings but that are clearly situated in the domain of perceptual inference. Specifically, we showed that neurons in the prearcuate gyrus can collectively represent abstract perceptual expectations before stimulus onset and confirmation biases during the evaluation of sensory evidence. These results suggest that neural mechanisms that perhaps originally evolved to select the best candidate action [37] over time acquired sufficient flexibility such that they can be repurposed to select the best candidate interpretation of the state of the environment in light of incoming sensory signals and prior experience.

Figurative language used to describe biased reasoning often invokes inertia. A biased person can be said to be entrenched in a position, immovable by evidence. In prefrontal cortex, we found a literal manifestation of the figurative relation between bias and inertia. When the neurally decoded decision variable covered less distance, monkeys gave a more biased account of the state of the sensory environment. This finding establishes a direct link between Bayesian computation and the dynamic evolution of neural activity during decision-formation. We suggest that this link exists because the brain’s implementation of Bayesian computation in our task resembles an approximate hierarchical inference process in which evidence is integrated until a bound is reached. It may seem surprising that estimating a static aspect of the environment involves temporal integration. However, noise in neural representations limits the signal-to-noise ratio of momentary sensory messages communicated by sensory cortex [30, 38, 39], and thus the quality of perceptual estimates informed by these messages [40]. Temporal integration averages out some of this noise, thereby improving the quality of perceptual estimates [39, 41, 42, 43, 44]. Under an approximate hierarchical inference process, prior expectations change the initial starting point and drift of the accumulator [35]. Using model simulations, we found that this process explains several key features of our data. First, it results in decisions that are biased for ambiguous as well as non-ambiguous stimulus orientations. Second, lowering stimulus contrast jointly reduces the slope of the psychometric function and increases the decision bias. Third, under this process, shorter excursions of the decision variable are associated with more biased and less sensitive decisions. We speculate that the neural implementation of Bayesian computation we have revealed is sufficiently general to support a wide range of perceptual and cognitive estimation tasks.

In this work, we provided a detailed description of the temporal evolution of population activity in prefrontal cortex during perceptual inference and we identified a principled computational process that accounts for these observations. It is natural to ask how neural circuits realize this computational logic. Specifically, how can hardwired circuits provided with a response mapping cue, a stimulus prior cue, and a snippet of noisy sensory measurements realize approximate hierarchical inference in a representational space that is orthogonal to the action selection space? We speculate that a recurrent neural network organized as an attractor network may be a fruitful starting place to address this question [45, 46, 47, 48]. Our analysis has provided a rich set of empirical constraints that should prove helpful to distinguish among candidate network organizations and training regimes and thus represents an important step towards uncovering the neurobiological basis of perception and cognition.

## METHODS

### Experimental methods and behavioral analysis

All analysis and simulation code for this work are included in a public github repository associated with this manuscript ^1^. The experimental methods are described in detail in ref. [3]. In brief, the experiments were performed on two adult male macaque monkeys (*Macaca mulatta*). The animals were trained to perform a memory-guided saccade task and an orientation discrimination task. All training, surgery, and recording procedures conformed to the National Institute of Health Guide for the Care and Use of Laboratory Animals and were approved by The University of Texas at Austin Institutional Animal Care and Use Committee. We first used a variation of the classical memory-guided saccade task [49] to identify recording sites where neurons exhibited neural activity indicative of an upcoming eye movement. Following identification of a suitable recording site (Extended Data Fig 1), we conducted several additional orientation discrimination training sessions with one choice target placed within the estimated response field location and one on the opposite site of the fixation mark. Once psychophysical performance reached a high level, physiological data collection began. The orientation-discrimination task involved two distinct prior contexts, associated with differently skewed distributions of stimulus orientation (see Fig. 1c). Blocks of both priors alternated randomly (80 trials per block). Stimuli were seven drifting gratings evenly spaced over a small range of orientation, tailored to each monkey’s orientation sensitivity. Vertically oriented stimuli received random feedback. Stimuli were presented at either high or low contrast (Michelson contrast: 100% or 4%). Blocks of high and low contrast stimuli alternated randomly (trials per block: monkey F = 100, monkey J = 80). We measured observers’ choice bias for ambiguous stimuli by computing the log odds of a “clockwise” choice under each prior context (Fig. 1d). We measured observers’ behavioral capability to discriminate stimulus orientation by fitting the relationship between stimulus orientation and probability of a “clockwise” choice with a psychometric function consisting of a lapse rate and a cumulative Gaussian function. Model parameters were optimized by maximizing the likelihood over the observed data, assuming responses arise from a Bernouilli process. Each recording session was analyzed independently. For the analysis documented in Fig. 1h, we fit one psychometric function per stimulus prior and contrast level. Both prior conditions shared the same sensitivity parameter, resulting in two psychometric functions with identical slope. We defined decision bias as the difference between the means of both cumulative Gaussians (i.e., the magnitude of the horizontal displacement of both psychometric functions). Error bars of model-based statistics are based on a 100-fold non-parametric bootstrap of the behavioral data.

### Simulated observer models

We investigated the choice behavior under an ideal Bayesian and a Maximum Likelihood inference strategy (Fig. 1e,f). We used two context-specific stimulus distributions that matched those used in the animal experiments. Each trial, the model observers were presented with a noisy sensory measurement, with the noise modeled as a sample from a zero mean Gaussian distribution. This sensory measurement informed the likelihood function, computed as a Gaussian probability density evaluated at all possible stimulus values, using a standard deviation that matched the strength of the sensory noise. For the Maximum Likelihood inference strategy, we selected the mode of the likelihood function as stimulus estimate. For the Ideal Bayesian inference strategy, we first used Bayes’ rule to compute the posterior probability over all possible stimulus values. The perceptual decision reflected whether a clockwise stimulus orientation was the most likely interpretation (meaning that the cumulative conditional posterior probability exceeded 50%).

### Electrophysiological recordings and decision variable analysis

As described in ref. [3], we conducted 13 successful recordings from monkey F and 16 from monkey J, using linear electrode arrays (average number of trials per session, monkey J = 3,171; monkey F = 1,593). We positioned the linear arrays so that they roughly spanned the cortical sheet and removed them after each recording session. To extract responses of individual units, we performed offline spike sorting. Given that the electrode’s position could not be optimized for all contact sites, most of our units likely consist of multi-neuron clusters. All units whose mean firing rate during the task exceeded 3 ips were included in the analysis. For each trial, we obtained moment-to-moment measurements of the decision variable by projecting 50 ms bins of population activity onto a linear decoder optimized to distinguish the activity patterns associated with both choice options (“left” vs “right” choices for the motor DV, and “clockwise” vs “counterclockwise” choices for the categorical DV, respectively). Specifically, we first individually z-scored each unit’s spike counts within every time bin. We then used these z-scored responses to estimate the set of linear weights, **w** = (*w*_1_, …, *w*_*n*_), that best separate the choice-conditioned z-scored response patterns, assuming a multivariate Gaussian response distribution:

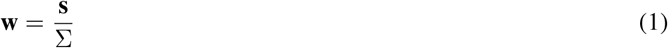

where **s** is the mean difference of the choice-conditioned z-scored responses and Σ is the covariance matrix of the z-scored responses. The decoder weights are calculated from observed trials. To avoid double-dipping, we excluded the trial under consideration from the calculation and solely used all other trials to estimate the weights. This way, we obtained “cross-validated” DV estimates for each time bin:

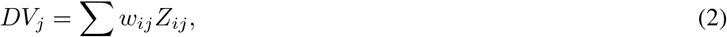

where *w*_*ij*_ and *Z*_*ij*_ are the weight and z-scored response of unit *i* on trial *j* for a given time bin. The symbols in Fig. 2a show example single trial DV trajectories.

For each trial, we fit a model with smoothly evolving DV-trajectories to the raw DV values (lines in Fig. 2a). This model has five free parameters: one captures an initial offset in the DV, one specifies the dynamic range of the DV trajectory, one controls the speed of the rise, one the time point at which half of the rise is completed, and one captures the decay in strength that follows the peak of the trajectory. The rising part of the trajectory follows a cumulative Gaussian profile, while the decay follows an exponential profile that begins at the time at which the cumulative Gaussian reaches the 99.38th percentile. We fit this model to the data by minimizing the sum of the squared error of each trial’s DV trajectory (correlations between model trajectory and raw DV trajectory are shown in Fig. 2b). We used these model trajectories to estimate, among other things, the peak DV value for each trial. To evaluate the usefulness of these estimates, we conduced a logistic regression analysis in which we predicted the choice behavior using two different sets of regressors. The first set comprised all experimentally controlled variables (stimulus prior, orientation, and contrast), the second set additionally included the peak DV value estimate. To compare the goodness of fit of both sets of regressors, we computed an estimator of prediction error based on information theory (Akaike’s information criterion, hereafter AIC). Specifically, assuming that the residuals under each model are distributed according to independent identical normal distributions:

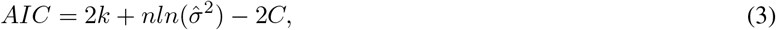

where *k* is the number of free parameters, *n* the number of data points, *C* a constant that only depends on the data, and 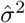 the maximum likelihood estimate for the variance of a model’s residuals distribution given by the residual sum of squares divided by the degree of freedom. Because only differences in AIC are meaningful, the constant *C* can be ignored when comparing models, yielding a statistic known as Δ*AIC* (shown in Fig. 2d).

We examined how the average peak DV value depended on the stimulus prior, orientation, and contrast (Fig. 2e-g). For each recording session, we quantified the effect of orientation with the slope of a linear regression line. We allowed these lines to have a prior-specific intercept, and conducted this analysis separately for the high and low contrast trials (example shown in Fig. 2e).

We studied the average temporal evolution of the DV trajectories for the zero-signal stimulus condition (Fig. 3a,b). To this end, we plotted the average DV around the putative time of choice commitment (–500 to –300 ms) computed from the model trajectory against the average value right before the onset of stimulus-driven activity (–800 to –600 ms). To quantify the growing influence of the prior expectation over the course of a trial, we calculated the first principle component of the resulting scatter plot (Fig. 3d). We evaluated the significance of the PC’s gradient by conducting a 1000-fold bootstrap-analysis and computing the 99.9% confidence interval from the resulting gradient estimate distribution.

### Simulated bounded accumulation process

We investigated the dynamic evolution of the decision variable in a bounded accumulation process and considered various scenarios. In all cases, the variable that was integrated was composed of one term representing the momentary sensory evidence, and one term representing the prior expectation. We modeled the momentary sensory evidence, *s*(*t*), at each time point *t* as a sample from a Gaussian distribution *s*(*t*) ∼ 𝒩 (*µ*_*θ*_, *σ*), with *µ*_*θ*_ proportional to the strength of the evidence (capturing the effects of stimulus orientation), and *σ* inversely proportional to the reliability of the evidence (capturing the effects of stimulus contrast). We reserved positive values to represent evidence for a clockwise stimulus orientation, and negative values evidence for a counterclockwise orientation. We modeled the prior expectation as a time-varying signal, *p*(*t*), that either only had energy at the first time point (capturing a pure stimulus expectation), or decayed to a lower but sustained level for the remaining time points (capturing an additional biased interpretation of sensory evidence). In addition, we introduced trial-by-trial variability to *p*(*t*) by multiplying this term with a scaling factor sampled from a log-normal distribution with a mean of one. Each trial consisted of 200 time steps, and each simulation consisted of 3,000 trials. When the simulated process contained decision bounds, the deliberation process was typically terminated early (at the time when one of the bounds was crossed). Fig. 3f illustrates an analysis in which we only included zero-signal stimuli and varied the strength of the prior-induced effects and the height of the bounds across simulations. Fig. 4a shows an analysis of the choice behavior for a simulation that included all of our experimental conditions as well as the effects of stimulus expectation, biased interpretation of sensory evidence, and terminating decision bounds.

### Dynamic range analysis

We investigated the relationship between the dynamic range of the DV excursion and behavioral signatures of decision bias and perceptual uncertainty. For each trial, we first computed the dynamic range of the DV by subtracting the DV’s initial value from the peak value. We then used a median split to divide all low-contrast (or high-contrast) trials of a single recording session (or model simulation) into a low- and high dynamic range group. Finally, we fit two sets of psychometric functions to the behavioral choices (one psychometric function per prior context, and one set of functions per dynamic range group). This yielded a total of four psychometric functions per session and contrast level (example for model simulation shown in Fig. 4c, and for real data in Fig. 4d).

To quantify the relationship between behavioral decision bias (Δ*β*) and the dynamic range of the neural DV, we used a metric, Δ*B*, defined as the surplus in prior-induced decision bias observed for the low dynamic range trials relative to the high dynamic range trials:

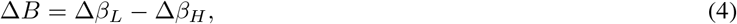

where Δ*β*_*L*_ is the prior-induced decision bias for the low dynamic range trials (specifically, the horizontal separation between the pair of psychometric functions fit to both prior contexts).

We devised an analogous metric, Δ*U*, to measure the association between perceptual uncertainty and DV dynamic range:

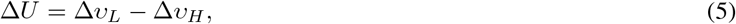

where Δ*υ*_*H*_ is the standard deviation of the cumulative Gaussian function that relates the proportion of clockwise choices to stimulus orientation for the high dynamic range trials.

## Acknowledgements

This work was supported by US National Institutes of Health grants T32 EY021462 (T.L., J.A.C.), EY032999 (R.L.T.G.), and National Science Foundation CAREER award #2146369 (R.L.T.G.).

## Author contributions

J.A.C. and R.L.T.G. conceived and designed the study. J.A.C. collected the data. T.L., J.A.C. and R.L.T.G. analyzed the data. T.L, J.A.C. and R.L.T.G. wrote the manuscript.

## Competing Interests

The authors declare no competing interests.

**Extended Data Figure 1.**
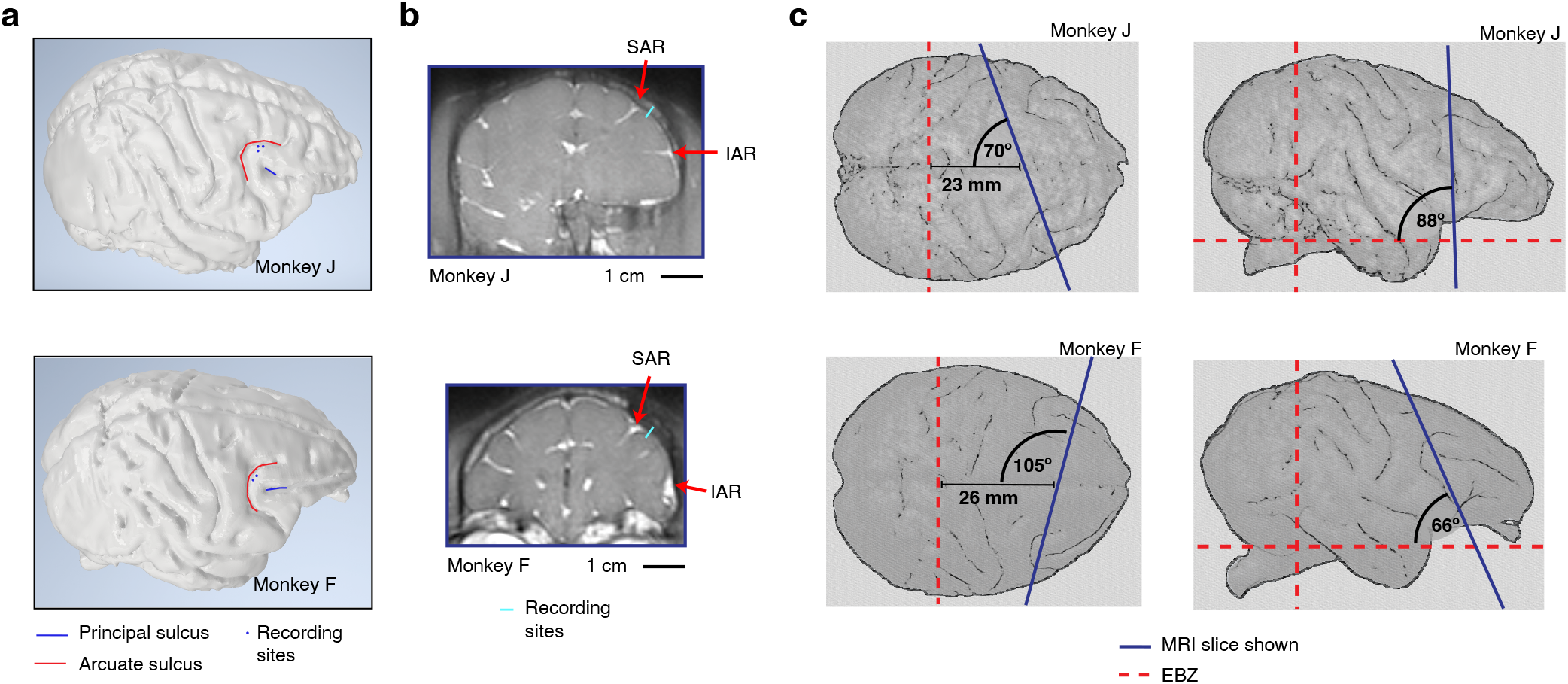
Location of prearcuate gyrus recordings for monkey J and F (top vs bottom). (**a**) 3D reconstruction of the brain based on a structural MRI scan obtained before chamber and post implants. The location of the recording sites are marked by blue dots. (**b**) Structural MRI scan illustrating the approximate recording site. The arrows indicate the superior arcuate sulcus (SAR) and inferior arcuate sulcus (IAR). (**c**) Top and side view (left vs right) of the MRI slice shown in panel **b**.

**Extended Data Figure 2.**
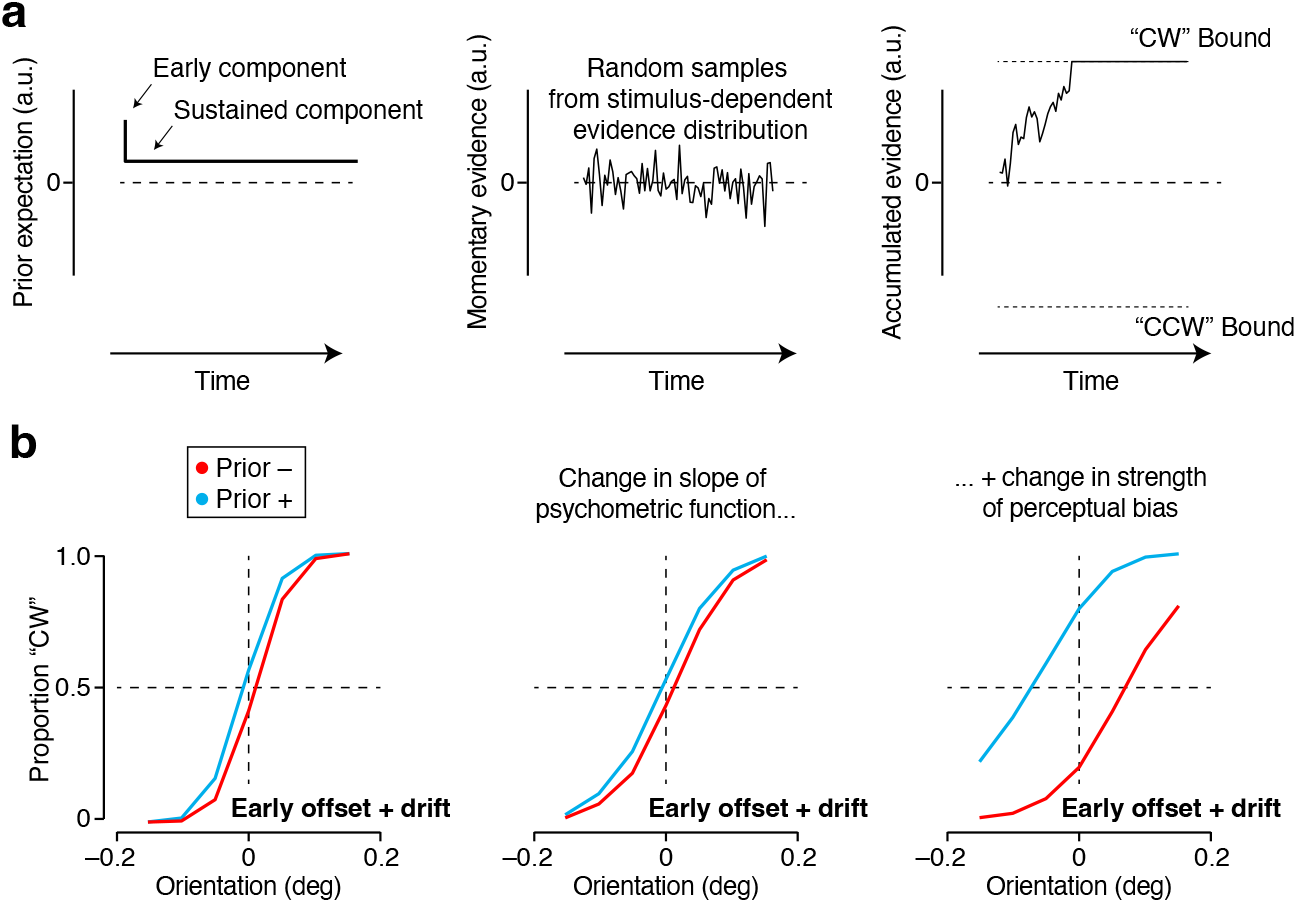
Simulated bounded accumulation process. (**a**) (left) The simulated bounded accumulation process contained three critical ingredients. First, a prior expectation, modeled as a time-varying signal with an early and late component. (middle) Second, a sensory input, modeled as a time-varying signal composed of random samples drawn from a Gaussian distribution. The mean of the distribution reflects stimulus orientation, the spread of the distribution stimulus contrast. (right) And third, two fixed bounds that terminate the accumulation process when crossed. (**b**) Under this process, prior expectations produce a robust horizontal shift of the psychometric function. The model parameters control the slope of the psychometric function and the magnitude of the decision bias.

**Extended Data Figure 3.**
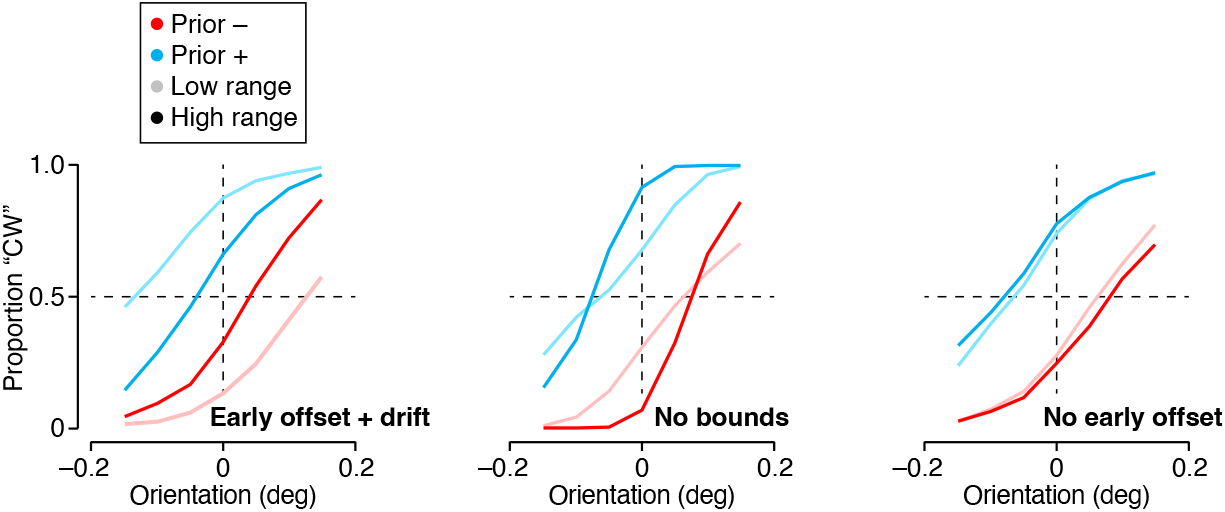
The relation between DV dynamic range and decision bias under different model variants. (Left) Median-split analysis of simulated choice behavior under two different prior expectations (blue vs red). The simulated process contained a prior-induced early offset and drift and terminating decision bounds. Trials with a smaller dynamic range yield more biased decisions (dark vs light colors). (Middle) Same analysis for an unbounded evidence accumulation process. Trials with a smaller dynamic range yield less sensitive decisions, as evidenced by the difference in the slope of the dark vs light psychometric functions. However, note that the decision bias, indicated by the horizontal separation of the midpoint of the red and blue psychometric function, does not depend on the dynamic range. (right) Same analysis for a bounded accumulation process in which the prior expectation does not induce an early offset, but only a drift.

## Footnotes

1 Project Github Repository link: https://github.com/andrefrancois22/Bayesian_Inference_PAG

## References

[1] Nikos K Logothetis and Jeffrey D Schall. Neuronal correlates of subjective visual perception. Science, 245(4919):761–763, 1989.

[2] Michael N Shadlen and William T Newsome. Neural basis of a perceptual decision in the parietal cortex (area lip) of the rhesus monkey. Journal of neurophysiology, 86(4):1916–1936, 2001.

[3] Julie A Charlton and Robbe L T Goris. Abstract deliberation by visuomotor neurons in prefrontal cortex. Nature Neuroscience, 27:1167–1175, 2024.

[4] Pascal Mamassian and Ross Goutcher. Prior knowledge on the illumination position. Cognition, 81(1):B1–B9, 2001.

[5] Wilson S. Geisler, J. S. Perry, Boaz J. Super, and D. P. Gallogly. Edge co-occurrence in natural images predicts contour grouping performance. 2001.

[6] Y. Weiss, E. P. Simoncelli, and E. H. Adelson. Motion illusions as optimal percepts. Nature Neuroscience, 5:598–604, 2002.

[7] Alan A Stocker and Eero P Simoncelli. Noise characteristics and prior expectations in human visual speed perception. Nature neuroscience, 9(4):578–585, 2006.

[8] Wendy J. Adams, Erich W. Graf, and Marc O. Ernst. Experience can change the ‘light-from-above’ prior. Nature Neuroscience, 7:1057–1058, 2004.

[9] David C. Knill. Learning Bayesian priors for depth perception. Journal of Vision, 7(8):13–13, 06 2007.

[10] Mehrdad Jazayeri and Michael N Shadlen. Temporal context calibrates interval timing. Nature neuroscience, 13(8):1020–1026, 2010.

[11] Julie A Charlton, Wiktor F Młynarski, Yoon H Bai, Ann M Hermundstad, and Robbe LT Goris. Environmental dynamics shape perceptual decision bias. PLOS Computational Biology, 19:e1011104, 2023.

[12] H. von Helmholtz. Handbuch der physiologischen Optik, volume III. Leopold Voss, 1867.

[13] D. Knill and A. Pouget. The bayesian brain: the role of uncertainty in neural coding and computation. Trends in Neurosciences, 27(12):712–719, ec 2004.

[14] A. R. Girshick, Landy M. S., and E. P. Simoncelli. Cardinal rules: visual orientation perception reflects knowledge of environmental statistics. Nature Neuroscience, 14:926–932, 2011.

[15] Christopher Summerfield and Floris P De Lange. Expectation in perceptual decision making: neural and computational mechanisms. Nature Reviews Neuroscience, 15(11):745–756, 2014.

[16] Wiktor F Mlynarski and Ann M. Hermundstad. Adaptive coding for dynamic sensory inference. Elife, 7:e32055, 2018.

[17] Michael Hahn and Xue-Xin Wei. A unifying theory explains seemingly contradictory biases in perceptual estimation. Nature Neuroscience, 27(4):793–804, 2024.

[18] Vinod Rao, Gregory C DeAngelis, and Lawrence H Snyder. Neural correlates of prior expectations of motion in the lateral intraparietal and middle temporal areas. Journal of Neuroscience, 32(29):10063–10074, 2012.

[19] Michele A Basso and Robert H Wurtz. Modulation of neuronal activity in superior colliculus by changes in target probability. Journal of Neuroscience, 18(18):7519–7534, 1998.

[20] Timothy D. Hanks, Mark E. Mazurek, Roozbeh Kiani, Elisabeth Hopp, and Michael N. Shadlen. Elapsed decision time affects the weighting of prior probability in a perceptual decision task. The journal of neuroscience, 31(17):6339–6352, 2011.

[21] Timothy R Darlington, Jeffrey M Beck, and Stephen G Lisberger. Neural implementation of bayesian inference in a sensorimotor behavior. Nature Neuroscience, 21(10):1442–1451, 2018.

[22] Hansem Sohn, Devika Narain, Nicolas Meirhaeghe, and Mehrdad Jazayeri. Bayesian computation through cortical latent dynamics. Neuron, 103(5):934–947, 2019.

[23] Vincent P Ferrera, Marianna Yanike, and Cassanello Carlos. Frontal eye field neurons signal changes in decision criteria. Nature Neuroscience, 12:1458–1462, 2009.

[24] Paul Cisek and John F Kalaska. Neural mechanisms for interacting with a world full of action choices. Annual review of neuroscience, 33:269–298, 2010.

[25] Jeffrey D Schall. Visuomotor areas of the frontal lobe. In Extrastriate cortex in primates, pages 527–638. Springer, 1997.

[26] Earl K Miller and Jonathan D Cohen. An integrative theory of prefrontal cortex function. Annual review of neuroscience, 24(1):167–202, 2001.

[27] Roozbeh Kiani, Christopher J Cueva, John B Reppas, and William T Newsome. Dynamics of neural population responses in prefrontal cortex indicate changes of mind on single trials. Current Biology, 24(13):1542–1547, 2014.

[28] Valerio Mante, David Sussillo, Krishna V Shenoy, and William T Newsome. Context-dependent computation by recurrent dynamics in prefrontal cortex. nature, 503(7474):78–84, 2013.

[29] W. P. Tanner and J. A. Swets. A decision-making theory of visual detection. Psychological Review, 61:401–409, 1954.

[30] P Heggelund and Albus K. Response variability and orientation discrimination of single cells in striate cortex of cat. Exp Brain Res., 32(2):197–211, 1978.

[31] Michael N. Shadlen, KH Britten, WT Newsome, and JA Movshon. A Computational Analysis of the Relationship between Neuronal and Behavioral Responses to Visual Motion. Journal of Neuroscience, 76(4):1486–1510, 1996.

[32] Robbe L T Goris, J. Anthony Movshon, and Eero P. Simoncelli. Partitioning neuronal variability. Nature Neuroscience, 17(6):858–865, 2014.

[33] Diogo Peixoto, Jessica R Verhein, Roozbeh Kiani, Jonathan C Kao, Paul Nuyujukian, Chandramouli Chandrasekaran, Julian Brown, Sania Fong, Stephen I Ryu, Krishna V Shenoy, and William T Newsome. Decoding and perturbing decision states in real time. Nature, 591(7851):604–609, 2021.

[34] Gabe Stine, Eric M Trautmann, Danique Jeurissen, and Michael N Shadlen. A neural mechanism for terminating decisions. Neuron, 2023.

[35] Richard Lange, Ankani Chattoraj, Jeffrey Beck, Jacob Yates, and Ralf M Haefner. A confirmation bias in perceptual decision-making due to hierarchical approximate inference. PLoS Comput Biol, 17(11), 2021.

[36] Trinity B Crapse, Hakwan Lau, and Michele A Basso. A role for the superior colliculus in decision criteria. Neuron, 97(1):181–194, 2018.

[37] Paul Cisek. Cortical mechanisms of action selection: the affordance competition hypothesis. Philosophical Transactions of the Royal Society B: Biological Sciences, 362(1485):1585–1599, 2007.

[38] David J Tolhurst, J Anthony Movshon, and Andrew F Dean. The statistical reliability of signals in single neurons in cat and monkey visual cortex. Vision research, 23(8):775–785, 1983.

[39] Rufin Vogels and G A Orban. How well do response changes of striate neurons signal differences in orientation: A study in the discriminating monkey. Jounal of Neuroscience, 10(11):3543–3558, 1990.

[40] Zoe Boundy-Singer, Corey M Ziemba, O J H Henaff, and Robbe L T Goris. How does v1 population activity inform perceptual certainty? Jounal of Vision, 24(6):1–17, 2024.

[41] Takanori Uka and Gregory C DeAngelis. Contribution of middle temporal area to coarse depth discrimination: comparison of neuronal and psychophysical sensitivity. Jounal of Neuroscience, 23(8):3515–3530, 2003.

[42] Leslie C Osborne, William Bialek, and Stephen G Lisberger. Time course of information about motion direction in visual area mt of macaque monkeys. Jounal of Neuroscience, 24(13):3210–3222, 2004.

[43] Yuzi Chen, Wilson Geisler, and Eyal Seidemann. Optimal temporal decoding of neural population responses in a reaction-time visual detection task. Jounal of Neurosphysiology, 99(3):1366–1379, 2008.

[44] Robbe LT Goris, Corey M Ziemba, J Anthony Movshon, and Eero P Simoncelli. Slow gain fluctuations limit benefits of temporal integration in visual cortex. Journal of vision, 18(8):8–8, 2018.

[45] Thomas Zhihao Luo,* Timothy Doyeon Kim,* Diksha Gupta, Adrian G Bondy, Charles D Kopec, Verity A Elliot, Brian DePasquale, and Carlos D Brody. Transitions in dynamical regime and neural mode underlie perceptual decision-making. Biorxiv preprint, 2024.

[46] Sebastian Bitzer, Jelle Bruineberg, and Stefan J Kiebel. A bayesian attractor model for perceptual decision making. PLoS Computational Biology, 11(8), 2015.

[47] Alexis Dubreuil, Adrian Valente, Manuel Beiran, Francesca Mastrogiuseppe, and Srdjan Ostojic. The role of population structure in computations through neural dynamics. Nature Neuroscience, pages 1–12, 2022.

[48] Mikhail Khona and Ila R Fiete. Attractor and integrator networks in the brain. Nature Reviews Neuroscience, 23:744–766, 2022.

[49] Shintaro Funahashi, Charles J Bruce, and Patricia S Goldman-Rakic. Mnemonic coding of visual space in the monkey’s dorsolateral prefrontal cortex. Journal of neurophysiology, 61(2):331–349, 1989.

